# Motivational control is implemented by a cingulo-prefrontal pathway

**DOI:** 10.1101/2025.02.11.637635

**Authors:** Clémence Gandaux, Jérôme Sallet, Céline Amiez, Delphine Autran-Clavagnier, Valentine Morel-Latour, Clément Goussi-Denjean, Vincent Fontanier, Pierre Misery, Camille Lamy, Frank Lamberton, Marina Lavigne, Eric J. Kremer, Charles.R. E. Wilson, Emmanuel Procyk

## Abstract

The neuronal connections between the midcingulate cortex (MCC) and the dorsolateral prefrontal cortex (dlPFC) are associated with multiple cognitive functions, including rapid and long-term adaptive processes. Here we show that DREADD-mediated activation of MCC to dlPFC projection in macaques led to increased engagement in a foraging task, but did not alter their trial-to-trial adaptive strategy. We conclude that a critical role for MCC-dlPFC pathway is in the temporally extended control of behaviour rather than in rapid adaptation.

---

Cortico-cortical dynamics and interactions within the frontal cortex are thought to drive adaptive cognitive functions. Cognitive flexibility is often associated with interactions between the midcingulate cortex (MCC) and dorsolateral prefrontal cortex (dlPFC), two regions of the primate brain associated with adaptive mechanisms implemented at different timescales ^1–3^. One classical example of “short term” adaptation to an event, is post-error adaptation ^4^ in which the MCC is proposed to monitor action outcomes and send information to the dlPFC about an immediate need to apply cognitive control and implement adaptation. Trial to trial signatures of behavioural adaptation include post-error slowing or win-stay lose-shift strategies. This process may be implemented via direct MCC-dlPFC connections, or alternatively via the locus coeruleus and noradrenergic control signals sent to the PFC ^5^. Neural dynamics in MCC and LPFC support such interactions ^6,7,8^. Other theoretical propositions focus on the role of MCC-dlPFC interaction in adaptation to information integrated over a much longer timescale. MCC neurons integrate outcome information over many trials^9^, and can provide an integrated signal to dlPFC to drive choices about when and how to engage in a task based on whether the current course of action provides acceptable outcomes. A clear example is the value signal used to decide whether to continue exploiting or shift toward exploring another patch during foraging ^10^. More generally, outcome integration can provide key information about whether to continue to engage in a task, hence contributing to aspects of motivation ^11^. This temporally-extended adaptation role contrasts with the trial-by-trial adaptation described above, and so would be expressed as rates of engagement in a task rather than single trial measures like post-outcome adaptation.

While the MCC and dlPFC may be associated with two distinct forms of adaptation, it is unclear whether or how they work together to achieve either or both. To date no causal observation has been reported. Understanding the role of the MCC-LPFC network, and the functions of the MCC in particular, is at the heart of current debates in cognitive neuroscience^12^. Whilst the effects of lesions in the individual regions MCC and dlPFC have been well studied ^9,13,14^, no experimental causal studies of their interactions exist. Yet a comprehensive understanding of network functions originates from understanding its interactions ^15^.

Here, we tested targeted modulation of the MCC to dlPFC pathway using network-specific chemogenetics in macaque monkeys performing an unconstrained homecage task that allows them to show evidence of adaptation at different timescales. We demonstrate for the first time that activating MCC to dlPFC projections does not perturb trial to trial adaptation but does alter the engagement in search behaviour.

We targeted the MCC-dlPFC pathway using *CAV-Cre*, a canine adenovirus (CAV-2) harbouring a Cre-recombinase expression cassette capable of retrograde transport to afferent regions, and *AAV hSyn-DIO-hM3Dq-mCherry*, an adeno-associated virus (AAV) harbouring a excitatory DREADD (***HM3Dq)*** flanked by inverted lox sequences (DIO) and an mCherry expression cassette ^13^. *CAV-Cre* was injected at multiple sites into posterior dlPFC and *AAV hSyn-DIO-hM3Dq-mCherry* was injected at multiple sites into MCC. We used this approach to exclusively express ***HM3Dq*** in cells projecting from MCC to dlPFC so that deschloroclozapine (DCZ)^16^ activated the MCC-dlPFC pathway (**Fig 1A, B**).

**Figure 1:**
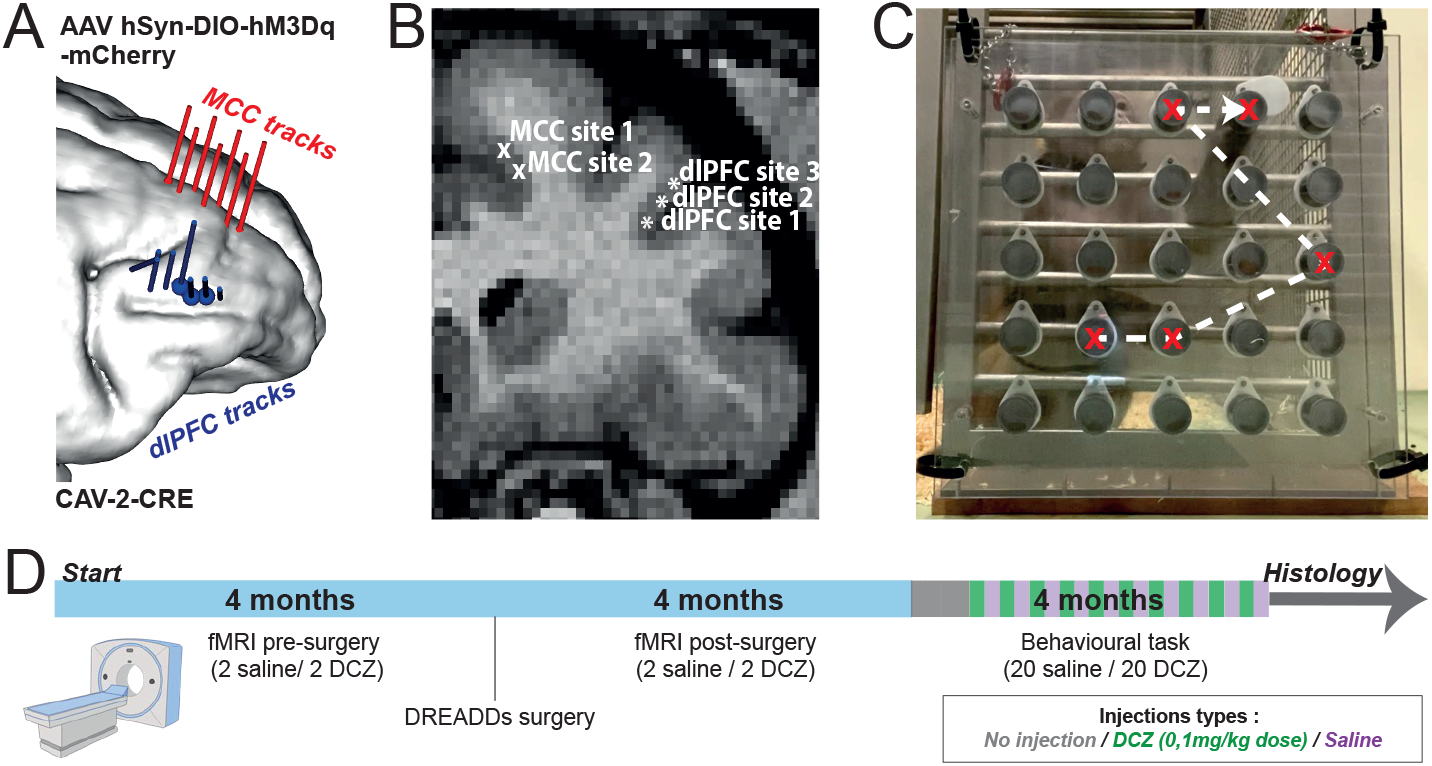
Injection sites and tracks and DCZ injection protocol. **A)** Tracks: Red lines represent injection tracks in the MCC of the *AAV hSyn-DIO-hM3Dq-mCherry* vector. Blue lines represent injection tracks in dlPFC of the *CAV-2-CRE* vector. **B)** Coronal MRI slice demonstrating injection sites in MCC and an injection track in dlPFC. **C)** Monkey performing a succession of sampled choices in the Spatial Foraging Task. **D)** Injection protocol for fMRI and behavioral studies: 4 fMRI scans were performed with DCZ (n=2) or saline (n=2) injection over 4 months prior to surgery. 3 Weeks after surgery the same protocols were performed. 8 months after surgery monkeys performed 20 sessions of the task with both DCZ and saline injections. In all our experiments the *DCZ dose was 0*.*1mg/kg*.

Animals performed a spatial foraging task in their homecage ^17,18^ in which they searched for 25 rewards in a patch containing 25 wells covered with a sliding transparent or opaque door depending on conditions. Each well contained a single reward. We analysed search strategies, choice types, and time spent engaged in the task (**Fig 1C**). In this task, monkeys with lesions in the lateral prefrontal cortex make more search errors than controls showing the task sensitivity to perturbation of the frontal cortex ^18^. This approach allowed us to identify adaptation at different timescales. We found that activating MCC to dlPFC projections does not perturb trial-to-trial adaptation, but rather alters the engagement in search behaviour.

Quantitative histological analyses showed MCC neurons expressing mCherry in both animals (**Fig. 2A**). These tagged neurons should only be those projecting to dlPFC, and are well localised in the injection sites in MCC (see histograms **Fig. 2E**). However, only a subset of MCC neurons monosynaptically project to the dlPFC. Using quantitative retrograde tracer studies by Markov et al (2014), we calculated the proportion of neurons in that specific pathway that should express hM3Dq. We estimated our mCherry labelling at 32% and 25% of MCC neurons projecting specifically to dlPFC in the two monkeys (details in Online Methods). As such, our method has targeted a significant proportion of the neurons in the pathway of interest, well above the 1% of neurons previously shown to be necessary for a behavioural effect in a cortical region ^19^. Moreover, the percentage of mCherry neurons that were in supragranular cortical layers (supragranular labelled neurons) was 35% and 50% for each monkey. These data thus suggest that the DCZ-mediated control of MCC neurons projecting to dlPFC acts mostly through feedback connections ^20^.

**Figure 2.**
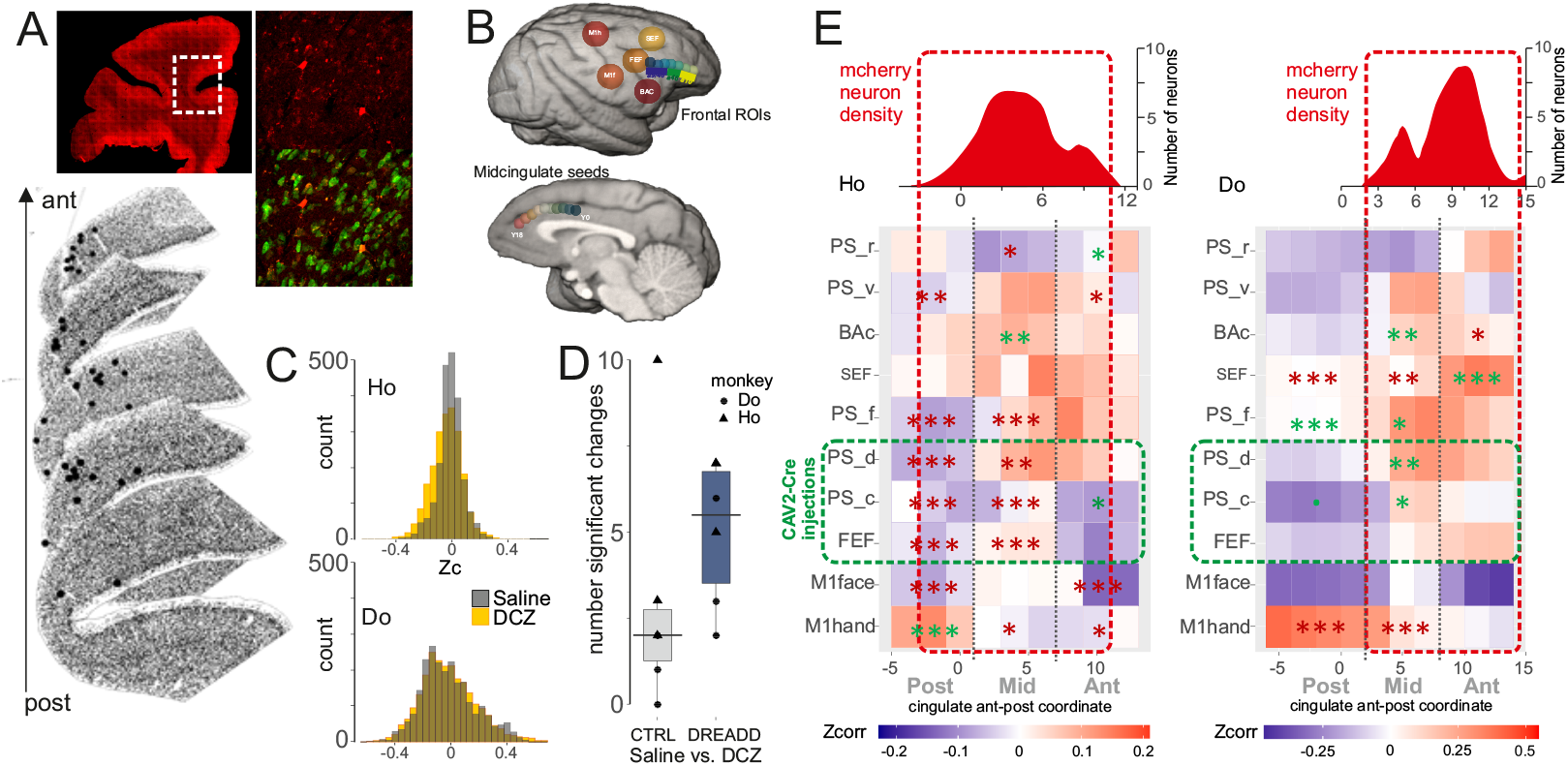
Anatomical and fMRI assessments of DREADDs-DCZ challenge on cingulate-to-frontal networks. **A**. Example of mCherry immunoreactive neurons in the fundus and dorsal bank of the cingulate sulcus (green: mature neurons NeuN). **B**. Positions of ROIs and seeds used for rs-fMRI analyses. **C**. Z-values across ***all*** ROI-Seeds couples in DCZ vs Control for each monkey (Ho and Do). Populations are not significantly different (*mixed linear model, fixed effects ‘condition’, random effects ‘session’ and ‘run’, p < 0*.*2 monkey H and p<0*.*9 monkey D*) **D**. Number of Zcorr values for couples of interest (see main text and panel E) that significantly differed between DCZ and Saline conditions in the pre-operative control phase (CTRL: no DREADD) and in the post-surgical DREADD phase. **E**. Top: red histograms show the density of mCherry immunoreactive neurons reported along the caudo-rostral axis of the cingulate sulcus for each monkey. Data are aligned at the level of the genu of the arcuate sulcus (0). Below: The heatmaps represent functional connectivity (Zc) observed in the saline condition between the cingulate dorsal bank (x axis) and the frontal ROIs (y axis). The stars represent statistical differences between saline and DCZ conditions for FC between ROIs and the 3 subdivisions of the cingulate (Post, Mid, Ant) (red stars reduction under DCZ; green stars increase under DCZ). Linear mixed effect models, *:p<0.05; **: p<0.01; ***: p<0.001. The Post-, Mid-, Ant-subdivisions were derived from data-driven clustering of seeds in previous work ^21^.

Using rs-fMRI under light isoflurane anaesthesia, we found that the overall functional connectivity (FC) measured between the entire cingulate sulcus (dorsal, fundus and ventral banks) and 10 regions of interest (ROIs) in the lateral frontal cortex (**Fig. 2B**, see supp. methods) was unchanged under DCZ compared to saline (**Fig. 2C**). However, when focussing on FC between the targeted dorsal bank of the cingulate with lateral frontal ROIs we found significant changes when comparing DCZ vs. saline sessions, particularly when compared to the same DCZ vs Saline contrast prior to surgery without hM3Dq (**Fig. 2D**). The studied rostro-caudal levels of the cingulate were functionally connected mostly to posterior parts of the principal sulcus (PS) (where CAV2-cre was injected), SEF and BA complex, and also with M1 face and hand regions (**Fig. 2E**, heatmap shows Z-correlations on the red-blue scale). These control levels of functional connectivity were modified in the DCZ+hM3Dq condition in both animals between dorsal cingulate regions (where mCherry+ neurons were mostly found, top histograms) and posterior regions of PS and FEF (where CAV-CRE was injected, green dashed box) (**Fig. 2E**, stars denote p<0.05). Within the different ROIs we mostly observed a decrease in FC under DCZ but not in all cases (**Fig. 2E**). Recent studies have shown that activating an inhibitory DREADD increases resting state FC ^22^.

Changes in functional connectivity under DCZ were found in lateral frontal areas beyond the zones targeted by CAV-Cre vector injections (putatively the projection zone of the labelled mCherry neurons). This demonstrates the engagement of a larger network - indirect connections or collateral branches of the infected cingulate neurons could be driving this effect.

In the spatial foraging task, activation with DCZ of MCC neurons projecting directly to dlPFC impacted the animals’ task engagement in the self-guided (opaque doors) condition but not in the visually-guided condition (transparent doors, **Fig. 3A,B**). Monkeys performed more trials overall in the self-guided condition, and continued searching for rewards for longer (for the less efficient monkey Ho, this also meant finding more rewards). DCZ alone without DREADDs has no impact on performance of this task (see supplementary data). This increased engagement meant animals showed more exploratory behaviour, and this in turn led to a longer delay before they repeated their choices under DCZ (**Fig. 3C**). Monkeys occasionally made short pauses in searching while remaining in front of the apparatus. The frequency of this behaviour was not impacted by DCZ (**Fig 3D**), but those pauses were later in the session in one animal with a trend the other (**Fig 3E**). These effects appear to be specific to the motivational engagement in the task. Note that this task is freely completed in a homecage with no constraints or food control applied.

**Figure 3.**
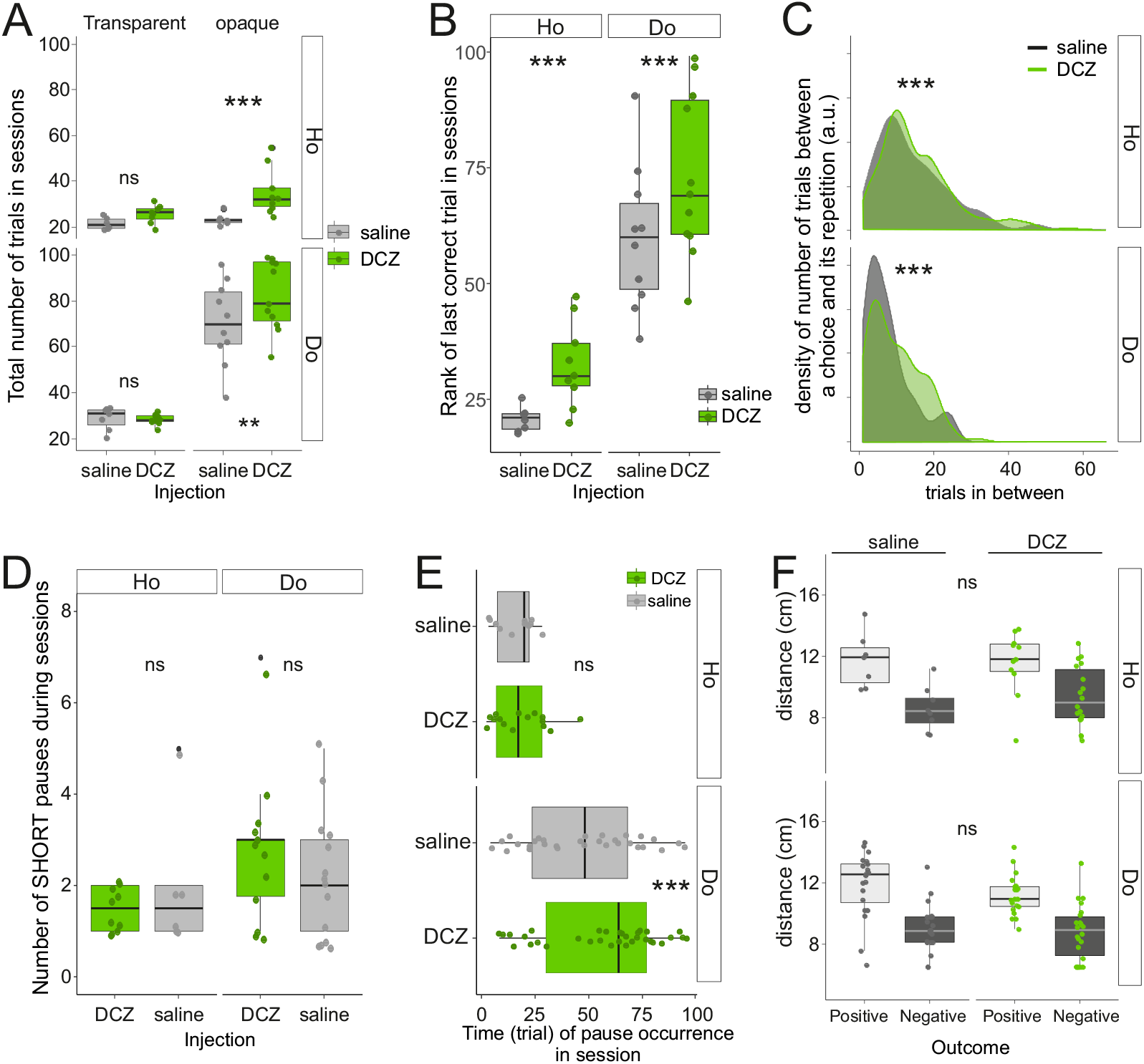
Behavioural effects of MCC-LPFC activation. **A**. Monkeys made more trials in the opaque condition under DCZ compared to saline. **B**. Monkeys gathered rewards over more trials under DCZ. **C**. The number of trials between a choice and its repetition was larger under DCZ compared to saline. **D**. Monkeys made as many short pauses in work in DCZ and sham sessions. **E**. Short pauses were made later in the session under DCZ in Do with a trend in Ho. **F**. In both monkeys the Euclidean distance to the next chosen well after a non-rewarded choice (negative outcome) was shorter than the distance after a rewarded choice. DCZ had no influence on this post-outcome effect. *: p < 0.05; **: p < 0.01; ***: p < 0.001.

By contrast, there was no impact of MCC-dlPFC pathway activation on the trial-to-trial adaptations occurring in this task. The monkeys showed a differential post-outcome adaptation in this task, by searching further away on the trial following reward compared to following no reward. This effect was not modified by DCZ injection (**Fig 3F**).

In conclusion, the targeted activation of the MCC-dlPFC pathway impacted sustained engagement in the task, and did not impact short term adaptation to outcomes. Our study provides the first conclusive data on the absence of a role of the MCC-LPFC pathway in trial-to-trial adaptation, a long debated subject with contradictory results when interventions or lesions are in one of the two regions ^9,13,23^. By contrast, we showed that the MCC-LPFC pathway contributes to sustained engagement in search or exploratory behaviours, supporting theoretical models in which MCC to LPFC interactions implement energizing behaviour, task maintenance, or a regulation of exploratory behaviours ^2,11,24,25^. This also suggests that outcome-related neural responses in MCC (a highly reproducible observation) might be more relevant for long term integration and value learning, rather than post-outcome or post-error trial-to-trial adaptation. Indeed, a neuropsychological study of medial to lateral frontal cortical interactions suggests an impairment in ERP measures of performance monitoring, but this impairment has no impact on trial-to-trial behaviour ^26^. Therefore, it can be argued that there is no consistent causal evidence of a trial-to-trial adaptation role for MCC in the literature. Our results are specific to the selective output of MCC to LPFC and collateral areas, rather than MCC function as a whole. One possibility is that MCC outcome processing contributes to post-outcome adaptation through other pathways which could be the target of future network specific interventions.

Globally, our study demonstrates the explanatory power of network-specific interventions in primates, allowing future studies to selectively modulate motivation or cognitive functions through network-specific targeting.

## Online methods

### Ethics and animals

Ethical permission was provided by “Comité d’Éthique Lyonnais pour les Neurosciences Expérimentales,” CELYNE, C2EA#42, under reference: APAFIS#14704_2018041318163087_v4. Monkey housing and care was done in accordance with the European Community Council Directive (2010) and the Weatherall report. Laboratory authorization was provided by the “Préfet de la Région Rhône-Alpes” and the “Directeur départemental de la protection des populations” under Permit Number: #B-690290402.

Two male adult rhesus macaque monkeys (Macaca mulatta) were included in the study: Monkey Ho., 15 years old, 14.00 kg; Monkey Do., 15 years old, 10.40 kg. Each animal was pair-housed with a female monkey. Both monkeys had been trained for a different cognitive task in a previous study ^1^.

Three other monkeys (Os, Yu and Vi), 2 males and 1 female rhesus macaques, were tested on the search task with and without DCZ im injection without viral injections.

### Chemogenetic

#### Surgical procedures and vector injections

We performed viral vector injections under aseptic conditions using previously published protocols for induction, anaesthesia and peri operative care ^2,3^. CAV-2 vector preparations were supplied through collaboration with the Kremer team at the Institut de Génétique Moléculaire de Montpellier, France, and the Plateforme de Vectorologie de Montpellier. AAV vector preparations were supplied commercially by AddGene.

For both monkeys, we performed a craniotomy over the dlPFC contralateral to the dominant hand of the animal (right hemisphere for Monkey Ho, left hemisphere for Monkey Do). We then opened and retracted the dura mater, to visualise the principal sulcus. Injection coordinates were chosen based on sites of maximal positive correlation of rsfMRI between the MCC and the dlPFC (see MRI procedures).

At each MCC site, we injected 1µL (1µL/min) of AAV5-hSyn-DIO-hM3Dq-mCherry. Each µL of vector contained 1*10^10^ particles of viral vector. We injected at multiple MCC locations separated by +1.5mm AP, along the dorsal bank and fundus of the cingulate sulcus at antero-posterior levels rostral to the genu of the arcuate sulcus (6 tracks for monkey H, 7 tracks for monkey D). At each location, we injected at two different depths within the cortex, separated by 1 to 1.5mm depending on the shape of the cingulate sulcus. The individual injection depths were MRI guided based on each individual animal’s local sulcal morphology, and depth calculations were made using Brainsight neuronavigation (Rogue Research, Canada). In total there were 12 injections in the MCC for both monkeys.

In parallel, in the same surgery and the same hemisphere, we made multiple injections into the dlPFC of CAV2-Cre. Each 1µl injection contained 3*10^9^ particles of viral vector. We injected at 7 dlPFC locations separated by +1.5mm AP along the dorsal bank of the principal sulcus and the anterior bank of arcuate sulcus / frontal eye field (FEF). At each location, we injected at two or three different depths separated by 1.5 to 3.5mm depending on the shape and depth of the dlPFC, and again guided by neuronavigation. In total there were 19 (Monkey H) and 20 (Monkey D) injections in the DLPFC.

Injections were made with 25µL Hamilton Syringes, with 22G & 30G needles. For MCC injections we used a syringe held in a standard stereotaxic arm and driven vertically. For dlPFC injections we used an injection arm supplied by Rogue Research as part of the Brainsight system, in order to angle injections in the axis of the principal sulcus derived from the animal’s MRI. For each injection track, we descended the needle to 0.5mm below the deepest injection site, waited for 10 minutes, then withdrew the needle to the injection site. After injecting at 1µL/min we waited for 1 min, before withdrawing the needle to the next depth. We waited 3 min before withdrawing the needle from the track after the final injection.

### DCZ preparation and injection protocols

For all DREADD activation and pre-operative injections of deschloroclozapine (DCZ), we used a dose of 0.1mg/kg. This dose has previously been shown not to induce off target effects in macaques ^4^. DCZ powder was put into solution in 1-2% in DMSO in advance and kept in individual aliquots for injection. On the day of injection, the DCZ was diluted in Q.S.P 0.1ml/kg of saline and the solution used within 24 hours and without further freezing. All injections were made I.M.

For behavioural experiments with the foraging task, we injected DCZ 30 minutes before testing monkeys. Intramuscularly injected DCZ has been shown to have significant behavioural effects within a few minutes in macaques, and to reach a significantly increased concentration in CSF within 30 minutes, an effect that lasts for at least 2 hours. Given that behavioural testing lasted 5 minutes, the testing can be considered to have occurred at the peak of effect of DCZ.

For MRI experiments (see MRI Procedures), we placed the anaesthetized animals in the scanner and acquired initial structural scans, before the I.M injections in situ. rsfMRI acquisitions reported in the results began 120 minutes post-sedation and 60 minutes after DCZ injection respectively.

### Behavioural Task

We tested the animals on a 25-option foraging task that aims to investigate spatial exploratory behaviour. This task was inspired by a previous paradigm designed to study working memory in primates ^5,6^; ^7^. The apparatus is made of a 400×400mm polycarbonate transparent panel in which 25 grey polycarbonate cylinders of 15mm diameter are inserted forming 25 wells. At the back of the setup, the cylinders are closed by a transparent polycarbonate panel. At the front, for each cylinder a door mounted on a pivot could be moved to access the content of the cylinders or wells. A food reward (e.g. a raisin) was placed in each well. The apparatus was placed outside of the home cage, at approximately 150mm, and held in position to the home cage by two safety hooks.

The animal had 1 min to start the task and had 5 min to interact with the device. Monkeys’ use of the apparatus was completely unconstrained during this time, but in each test case the optimal behaviour would be to retrieve all of the rewards, 1 from each well, without returning to previously explored wells, as they were never re-baited during testing. During the exploration, the animal was allowed to pause for a maximum of 1 min - beyond this delay, the session was stopped. Polycarbonate doors were either transparent or white opaque. The transparent doors permitted the monkeys to visually verify where the rewards were, whereas opaque doors required animals to use memory if they wished to avoid repetitive choices. The transparent condition was named ‘control’ and the opaque condition ‘test’ sessions.

All sessions were videotaped using a smartphone to allow offline behavioural analysis. Nature of the trials (correct, incorrect or missed trials) and position of the chosen well were obtained from the videos. Sessions of less than 15 trials were discarded.

To prevent analysis bias, data collection and data analysis were conducted by different researchers. The researcher coding the video was unaware of the nature of the sessions being analysed (saline, dreadds, or no injection).

Animals were introduced to the foraging task approximately 3 months following the vector injections. Data collection started approximately 5 months after the surgery.

### rs-fMRI

#### Acquisition

We performed 4 resting state fMRI scans pre-surgery and 4 post-surgery. In each phase (pre- and post-op) monkeys received injection of the DREADD activating ligand DCZ ^8^ (0.1mg/kg) for 2 scans, and monkeys received control vehicle injections for the other 2. This balanced design allowed us to test for non-specific effects of DCZ before DREADD injection ^4^, as well as the specific DREADD-driven effect. to determine targets for viral infections. Post operative scans were acquired 10 weeks after surgery.

Data were acquired on a 3T Siemens Magnetom Prisma MRI scanner (Siemens Healthineers, Erlangen, Germany). Prior to anaesthesia, monkeys were injected with glycopyrrolate, an anticholinergic agent that decreases salivary secretion (Robinul; 0.06mg/kg). Twenty minutes after, anaesthesia was induced with an intramuscular injection of tiletamine and zolazepam (Zoletil; 7 mg/kg). The animals were then intubated and ventilated with oxygen enriched air. Anaesthesia was maintained with 0.8% Isoflurane throughout the duration of the scan, with the aim of maintaining a stable but light plane of anaesthesia. Monkeys were placed in an MRI compatible stereotaxic frame (Kopf, CA, USA) in a sphinx position. During the scan, physiological parameters including heart rate and ventilation parameters (spO2 and CO2) were monitored. Body temperature was measured and maintained using warm-air circulating blankets. Resting-state fMRI acquisitions started 2 hours after anaesthesia induction and 60 min after DCZ or saline injections. The aim was to allow significant washout of the inducing agent prior to rs-fMRI acquisitions ^9^, and to ensure that DCZ was maximally present after an intramuscular injection ^8^. L11 Siemens loop coils were placed vertically on each side of the monkey’s head and 1 L7 Siemens was placed horizontally above the monkey’s head. A high resolution T1-weighted anatomical scan was first acquired (MPRAGE, 0.5mm3 isotropic voxels, 144 transverse slices, TR=3000ms, TI=1100ms, TE=3,62ms, 2 averages). Resting-state functional images were obtained in an ascending order with a T2*-weighted gradient echo planar images (EPI) sequence with the following parameters: TR=1900 ms, TE=30ms, flip angle = 75°, 28 transverse slices, isotropic voxel size: 1.7mm3. We collected 5 runs per session. With 400 volumes per run (12 min each) for a total of 16 000 volumes across the 2 animals and all sessions.

### Rs-fMRI data analysis

The preprocessing of resting-state scans was then performed with SPM 12. The first 5 volumes of each run were removed to allow for T1 equilibrium effects. First, we performed a slice timing correction using the time center of the volume as reference. The head motion correction was then applied using rigid body realignment. Then, images were skull-stripped using the bet tool from the FSL software (https://fsl.fmrib.ox.ac.uk/fsl/fslwiki/BET, Jenkinson et al. 2005). Using the SPM software, the segmentation of each brain of each session was performed on skull-stripped brains. To ensure optimized inter-session and inter-subject comparisons, both anatomical and functional images were then registered in a common atlas space, CHARM/SARM ^10,11^ (see https://afni.nimh.nih.gov/pub/dist/doc/htmldoc/nonhuman/macaque_tempatl/atlas_charm.html). Temporal filtering was then applied to extract the spontaneous slowly fluctuating brain activity (0.01–0.1Hz). Finally, linear regression was used to remove nuisance variables (the cerebrospinal fluid, white matter signals from the segmentation, and volumes containing artefacts as detected by the ART toolbox, https://www.nitrc.org/projects/artifact_detect/) and spatial smoothing with a 4-mm FWHM Gaussian kernel was applied to the output of the regression.

#### Seed Selection in the MCC

The seeds consisted of 2.5mm radius spheres positioned in the cingulate sulcus (covering both ventral and dorsal banks of the cingulate sulcus) in hemisphere right for monkey Ho and left for monkey Do, starting 10 mm posterior to the anterior commissure to the rostral end of the cingulate sulcus, and spaced from 2.5mm each, for a total of 11 seeds. (CgS1, CgS2, CgS3, CgS4, CgS5, CgS6, CgS7, CgS8, CgS9, CgS10 and CgS11). Given that, in macaques, CMAr is located about 10mm anterior to the genus of the arcuate, we anticipated that this subdivision would roughly correspond to CgS8 ^12^. Caudal seeds CgS1 to CgS7 correspond to Y values -5 to +10 (with AC at Y0) whereas CgS8 to CgS11 correspond to Y values +12.5 to +20. According to our previous meta-analysis aiming at identifying the location of the rostral cingulate motor area (CMAr) in macaques ^12^, the face motor area CMAr is located about 10mm anterior to ArcGen.

#### Selection of Region Of Interest (ROI) in the dorsolateral prefrontal cortex

For both monkeys we chose on a rostro-caudal axis areas 46 and 9/46 of the dlPFC. Area 9/46 occupies the caudal part of the principal sulcus, and area 46 occupies the principal sulcus between area 9/46 and area 10 ^13^.

#### Correlations between seeds and ROIs

For each hemisphere of each animal, Pearson correlation coefficients between the seeds with the various ROIs in the prefrontal cortex were computed and normalized using the Fisher’s r-to-z transform formula. The significant threshold at the individual subject level was set to Z = 0.1 (p < 0.05). These normalized correlation coefficients, which corresponded to the functional connectivity strength between each seed and each ROI in individual brains, were subsequently processed with R software for all the following analyses.

### Histology

At the end of the protocol, monkeys were sacrificed and perfused for histological analysis. Anesthesia was induced with an IM injection of ketamine (Imalgene 10mg/kg) and then the animal was euthanized by injection of pentobarbital (Euthasol Vet, 1ml/kg, IV). Brain tissue was then perfused transcardially with a 2.7% saline + 1% procaine, 4% PFA in 0.1M of phosphate buffer (pH=7.4) and fixed sequentially with 5%,10%, and 20% of glycerol.

Brains were cut frozen in coronal sections of 40μm thickness from the frontal lobe with a sliding microtome. We defined a region of interest (ROI) including the injection sites between 8.5 and 24mm and 9mm and 26mm after the rostral tip of the brain for monkey Do and monkey Ho respectively.

We used immuno-histo-fluorescence in 4° C stocked slices. Slices were washed in 0.1M PB. They were blocked and permeabilized in TNB-Tx0.4% composed of 0.1M PB, casein 0.05% (SIGMA), BSA 0.25% (SIGMA), and Triton® 0.4% (Alfa Aesar). Primary antibodies used were guinea pig anti-NeuN (266004 SYNAPTIC SYSTEM 1:2500) to mark all mature neurons, and rabbit Anti-RFP (PM005 CLINISCIENCES 1:1000) to mark mCherry tagged neurons that should also be expressing the DREADD receptors. They were incubated with TNB-Tx 0.4%. Before moving to secondary antibodies, slices were washed in PB-Tx 0.25%. Secondary antibodies were goat anti-guinea pig Alexa Fluor 488 (A-11073 THERMOFISHER 1:500) and donkey anti-rabbit Biot (711-065-152 JAKSON LABORATORIES 1:500) and they were incubated with TNB-Tx 0.4%. Amplification of donkey anti-rabbit Biot was carried out with streptavidin 555 in TNB-Tx (016-160-084 JAKSON LABORATORIES 1:500). The nuclei were marked with DAPI. The sections were immediately mounted from 0.1 M phosphate buffer (pH=7.4) solution onto 3% gelatin-coated slides. Sections were mounted with fluoromount.

We mounted 24 slices at a sampling frequency of 1/18. Mounted sections were scanned in a ZEISS Axioscan 7. Then they were analysed with Neurolucida 360 ® MBF bioscience software. mCherry neurons were sparse and were counted by hand, whereas NeuN counting of mature neurons used automatic counting within the software. To estimate the total number of mCherry labelled neurons we used a statistical model of interpolation based on the calculation of the mean standard deviation relation for the plotted cells, an approach derived from the retrograde tracer technique ^14^. Analyses were carried out in R. We estimated the proportion of MCC neurons expressing mCherry at 0.07% and 0.05% in the two monkeys.

However, only a subset of MCC neurons are monosynaptically projecting to the DLPFC, and so we aimed to estimate the proportion of these mCherry labelled neurons as a function of the total projection from MCC to dlPFC, but this total number is not obtainable without direct anatomical tracing that would have blocked marking of the mCherry labelled neurons. In order to overcome this limitation, we therefore used gold-standard anatomical tracing data for the size of the MCC-dlPFC projection, albeit from different monkeys. Using the Markov et al database, we counted the number of MCC neurons projecting to the dlPFC. Based on the detailed parcellation provided in the database, we considered a combination of 9/46d and 46d to be the equivalent extent of our dlPFC ROI for the labelled neurons. The area 24C was the equivalent of our MCC ROI. In their study they estimated the number of area 24c neurons projecting to these two different locations to n=4542. Using these high-quality data as baseline for the size of the projection, albeit derived from different macaques, we estimated we have labelled 32% and 25% of MCC neurons projecting to dlPFC in the two monkeys.

## SUPPLEMENTARY FIGURES

**Supplementary Figure 1.**
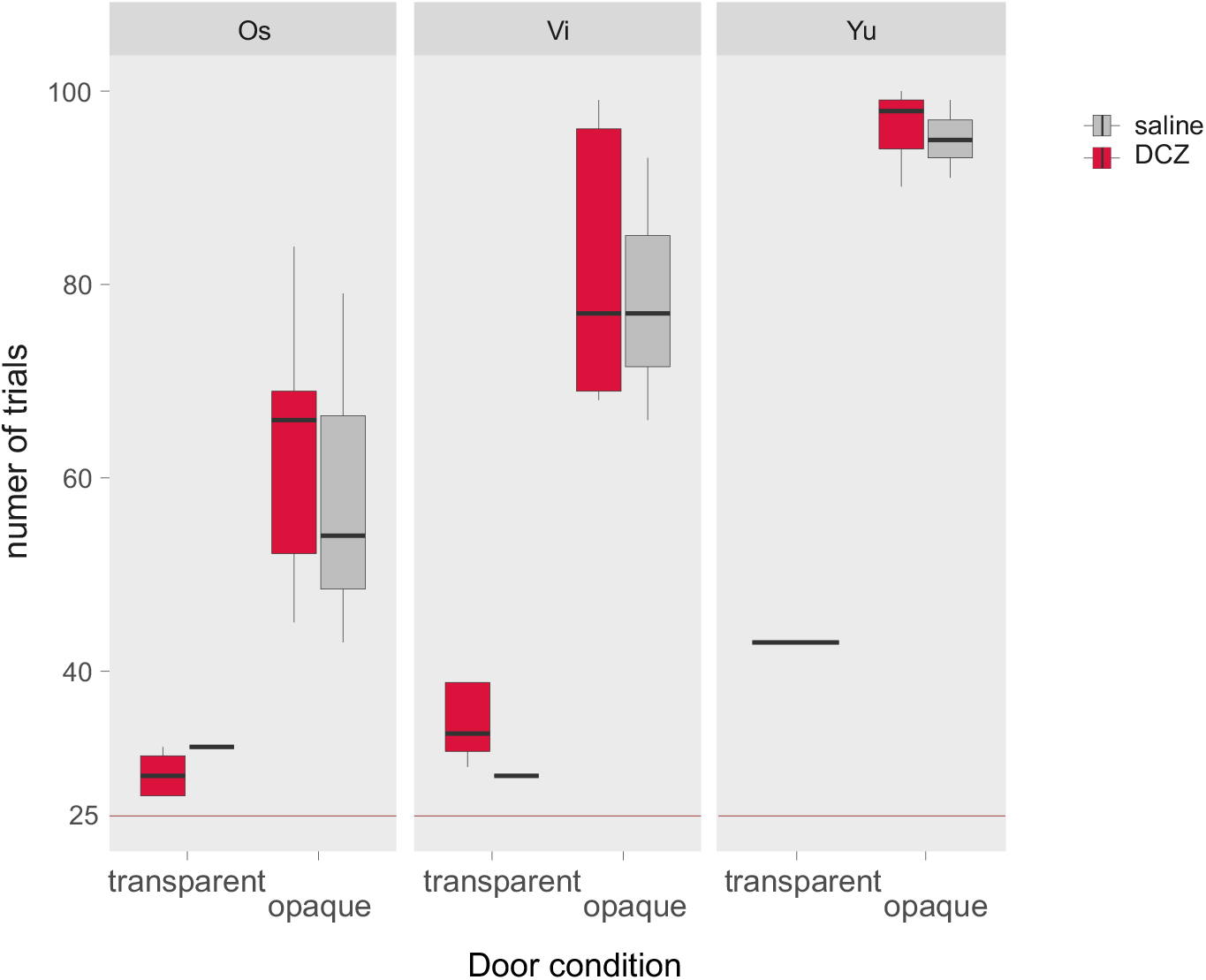
Testing DCZ injection on trial number with no DREADD. The figure shows the total numbers of trials for 3 monkeys under saline or DCZim injections in the foraging task, with transparent or opaque doors. No effect of injection was found (linear mixed effect model, fixed interaction terms: Injection x Condition, random effect: monkey). interaction: ns, Condition p<10-5, Injection ns).

